# Looking at the Full Picture: Utilizing Topic Modeling to Determine Disease-Associated Microbiome Communities

**DOI:** 10.1101/2023.07.21.549984

**Authors:** Rachel L. Shrode, Nicholas J. Ollberding, Ashutosh K. Mangalam

## Abstract

The microbiome is a complex micro-ecosystem that provides the host with pathogen defense, food metabolism, and other vital processes. Alterations of the microbiome (dysbiosis) have been linked with a number of diseases such as cancers, multiple sclerosis (MS), Alzheimer’s disease, etc. Generally, differential abundance testing between the healthy and patient groups is performed to identify important bacteria (enriched or depleted in one group). However, simply providing a singular species of bacteria to an individual lacking that species for health improvement has not been as successful as fecal matter transplant (FMT) therapy. Interestingly, FMT therapy transfers the entire gut microbiome of a healthy (or mixture of) individual to an individual with a disease. FMTs do, however, have limited success, possibly due to concerns that not all bacteria in the community may be responsible for the healthy phenotype. Therefore, it is important to identify the *community* of microorganisms linked to the health as well as the disease state of the host.

Here we applied topic modeling, a natural language processing tool, to assess latent interactions occurring among microbes; thus, providing a representation of the *community* of bacteria relevant to healthy vs. disease state. Specifically, we utilized our previously published data that studied the gut microbiome of patients with relapsing-remitting MS (RRMS), a neurodegenerative autoimmune disease that has been linked to a variety of factors, including a dysbiotic gut microbiome.

With topic modeling we identified communities of bacteria associated with RRMS, including genera previously discovered, but also other taxa that would have been overlooked simply with differential abundance testing. Our work shows that topic modeling can be a useful tool for analyzing the microbiome in dysbiosis and that it could be considered along with the commonly utilized differential abundance tests to better understand the role of the gut microbiome in health and disease.

**Author Summary:** Trillion of bacteria (microbiome) living in and on the human body play an important role in keeping us healthy and an alteration in their composition has been linked to multiple diseases such as cancers, multiple sclerosis (MS), and Alzheimer’s. Identifying specific bacteria for targeted therapies is crucial, however studying individual bacteria fails to capture their interactions within the microbial community. The relative success of fecal matter transplants (FMTs) from healthy individual(s) to patients and the failure of individual bacterial therapy suggests the importance of the microbiome *community* in health. Therefore, there is a need to develop tools to identify the communities of microbes making up the healthy and disease state microbiome. Here we applied topic modeling, a natural language processing tool, to identify microbial communities associated with relapsing-remitting MS (RRMS). Specifically, we show the advantage of topic modeling in identifying the bacterial community structure of RRMS patients, which includes previously reported bacteria linked to RRMS but also otherwise overlooked bacteria. These results reveal that integrating topic modeling with traditional approaches improves the understanding of the microbiome in RRMS and it could be employed with other diseases that are known to have an altered microbiome.

## Introduction

The microbiome is the collection of microorganisms that live in and on our body. Although, the microbiome includes bacteria, viruses, fungi, and phages, the majority of microbiome studies have been focused on bacteria. With regard to the bacterial microbiome, it has been established that there is a community structure where a number of different species from various bacterial phyla live together. Their composition is regulated by various host and microbe specific factors and in a steady state, they help to maintain homeostasis, keeping the host healthy. However, the alteration in the composition of the microbiome (dysbiosis), has been linked to a number of diseases including cancers, multiple sclerosis, Parkinson’s disease, Alzheimer’s disease, inflammatory bowel disease (IBD), and others [1–4]. In the majority of microbiome studies, the relative abundance of each individual microbe is compared one at a time between people with a particular disease and healthy controls. This type of analysis has provided several major findings on overly enriched or overly depleted microbes that are linked to disease. For example, *Fusobacterium nucleatum* with colorectal cancer [5] and *Clostridium difficile* with IBD [6].

These findings are helpful in each respective area of research however, providing a singular species of bacteria to an individual lacking that species for health improvement has not been as successful as fecal matter transplant (FMT) therapy. A fecal matter transplant (FMT), where the entire microbiome is provided, the recipient can see improvement of disease [7–9]. This reveals to us that the *community* of microorganisms is important to our health, and we should consider the structure of the community to better prevent, diagnose, and treat disease. FMTs do, however, have limited success possibly due to concerns that not all bacteria in the community may be responsible for the healthy phenotype. Therefore, there is a need for a method to identify communities within the healthy community.

In this work we aim to show the benefits of using the natural language processing (NLP) tool, topic modeling, in order to assess the *community* structure associated with diseases. Topic modeling is an unsupervised machine learning approach that assesses all the terms (bacteria) within documents (samples) and groups them into topics (communities) based on term similarities and patterns.

To do so, we utilized our previously published data on the gut microbiome composition of relapsing-remitting multiple sclerosis (RRMS) patients. RRMS is a neuroinflammatory autoimmune disease caused by genetic and environmental factors. The gut microbiome has emerged as a major environmental factor of interest in the development of RRMS as many studies have revealed that RRMS patients have a dysbiotic gut microbiome [10–18]. As previous studies have focused on individual microbial differences, we instead applied topic modeling to our RRMS gut microbiome data to assess the latent interactions occurring among microbes and their association with RRMS. Specifically, we used the Latent Dirichlet Allocation (LDA) model as it allows documents to have fractional membership across topics [19]. With topic modeling we in fact were able to confirm previously identified bacteria of interest linked with RRMS, but we additionally identified *communities* of bacteria, with otherwise overlooked bacteria, linked to RRMS. Therefore, we suggest topic modeling in addition to traditional approaches to better understand the microbiome of individuals with RRMS and other disease with dysbiotic microbiome communities.

## Results

### Data For Analysis

Our primary analysis was performed on the data from Chen et. al. 2016 [11] and validated with the data from Yadav et. al. 2022 [3]. For simplicity, each dataset is referred to by using the first author’s last name (e.g., Chen and Yadav). After data processing, we retained 175 genera in the Chen dataset and 160 in the Yadav validation dataset.

### Number and Similarity of Validated Community Types

Our cosine similarity analysis revealed 34 community-type associations with high correlations (> 0.80) between the Chen and Yadav topics (Fig. 1), highlighting similarities in the community types across datasets. These associations comprised 13 topics from the exploratory dataset and nine from the validation dataset that were also associated with RRMS versus HC status. In the Chen dataset, 10 of the 30 topics were enriched in samples obtained from the RRMS patients compared to controls (Fig 2). In the Yadav dataset, four of the 30 topics were associated with RRMS versus HC status, with three enriched in RRMS and one enriched in HC samples. The plots for all statistically significant topic associations can be found in Supplementary Figure 2.

**Fig 1:**
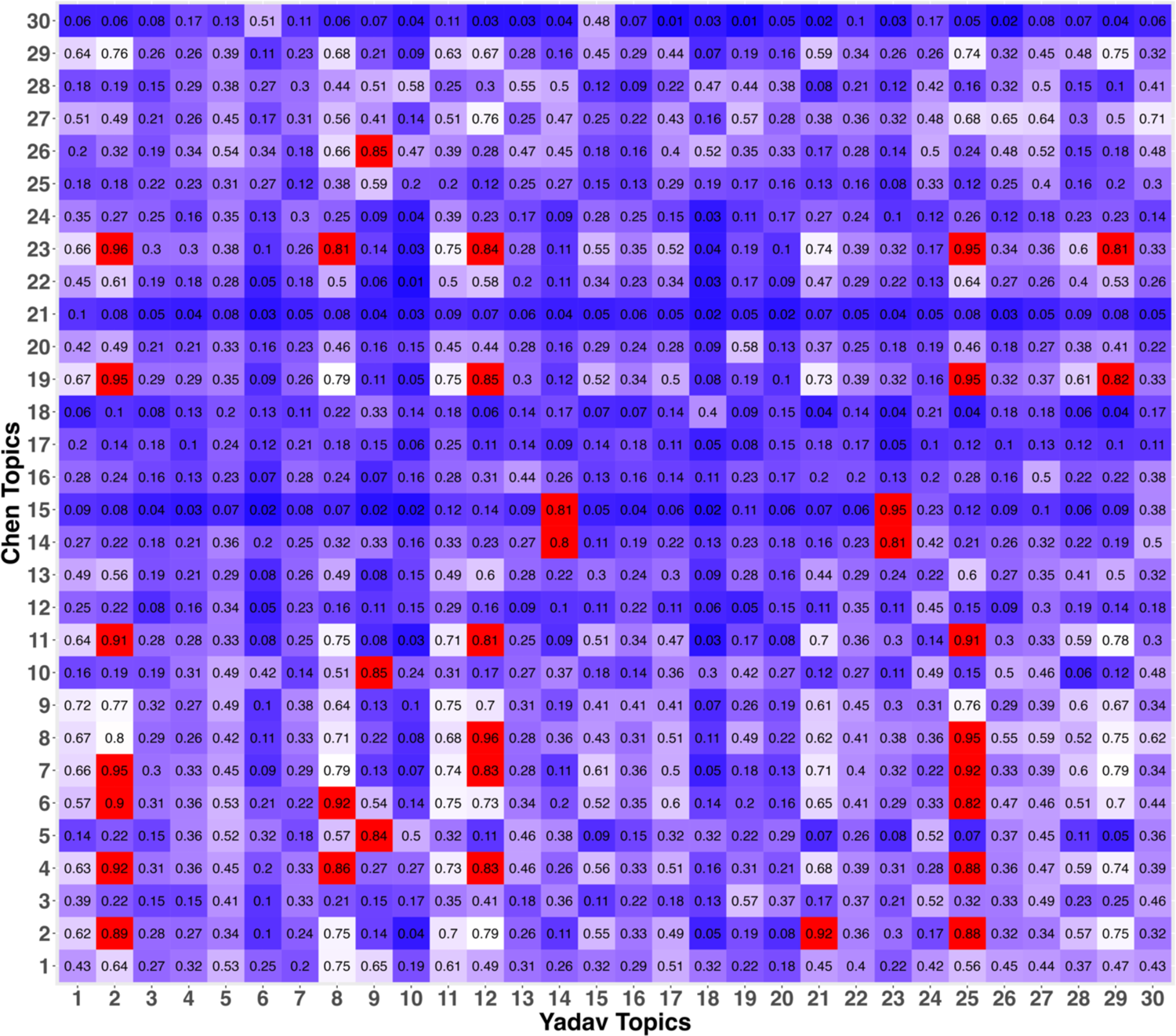
Community structure (topic) cosine similarity between Chen and Yadav datasets. A higher value reveals similar term assignment to topic. A value of 0.80 or greater was considered to reflect similar community types.

**Fig 2:**
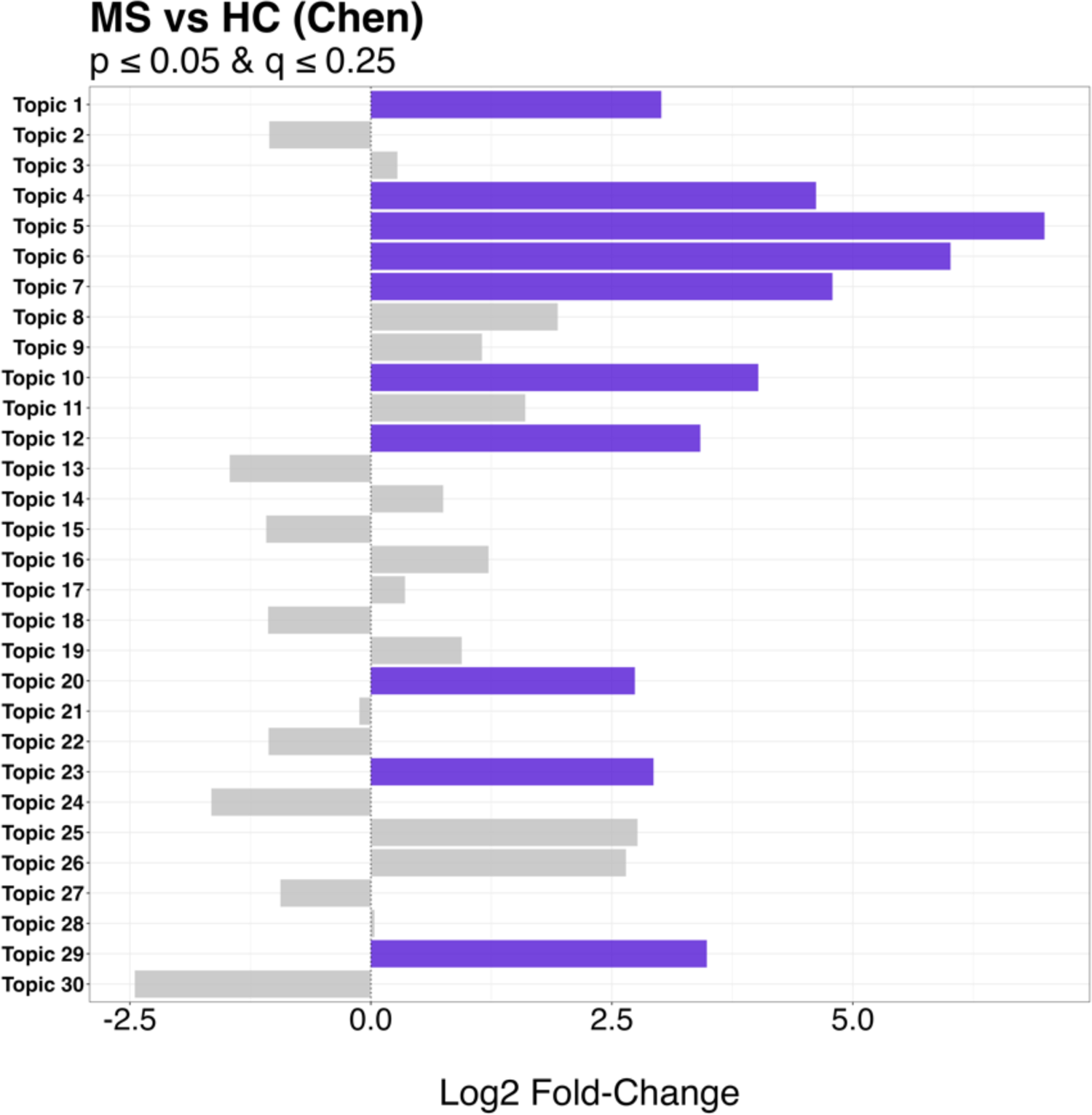
Differentially abundant community types. Statistically significant topics assigned in Chen dataset highlighted in purple.

Specifically, of the significant community types found in the Chen data, five were validated based on having high cosine values (> 0.80) to topics derived independently from the Yadav validation data. All these topics were significantly (p ≤ 0.05 and q ≤ 0.25) enriched in RRMS patients compared to HC. In detail, Chen Topic 4, Chen Topic 6, and Chen Topic 23 were similar to Yadav Topic 8 (cosine = 0.92, 0.86, 0.81, respectively). We will refer to this validated community as Community Type A (Fig. 3.a). Chen Topic 5 and Chen Topic 10 were similar to Yadav Topic 9 (cosine = 0.84, 0.85, respectively). We will refer to this validated community as Community Type B (Fig. 3.b). For an overview of the topics making up these Community Types see Fig. 4.a and 4.b.

**Fig 3a:**
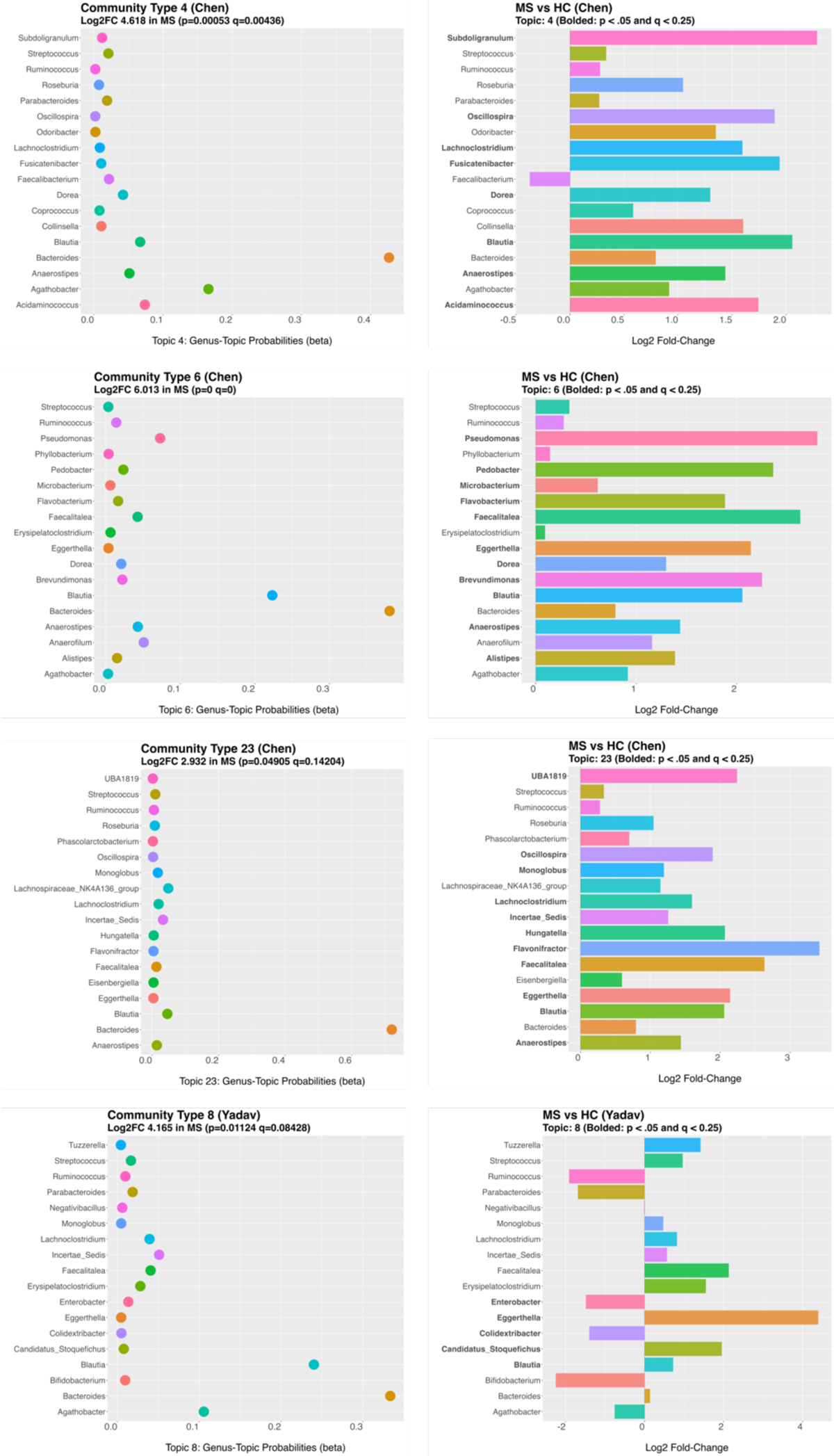
MS gut microbiome Community Type A found in Chen and validated in Yadav. Specifically, communities Chen Topic 4, Chen Topic 6, Chen Topic 23, and Yadav Topic 8.

**Fig 3b:**
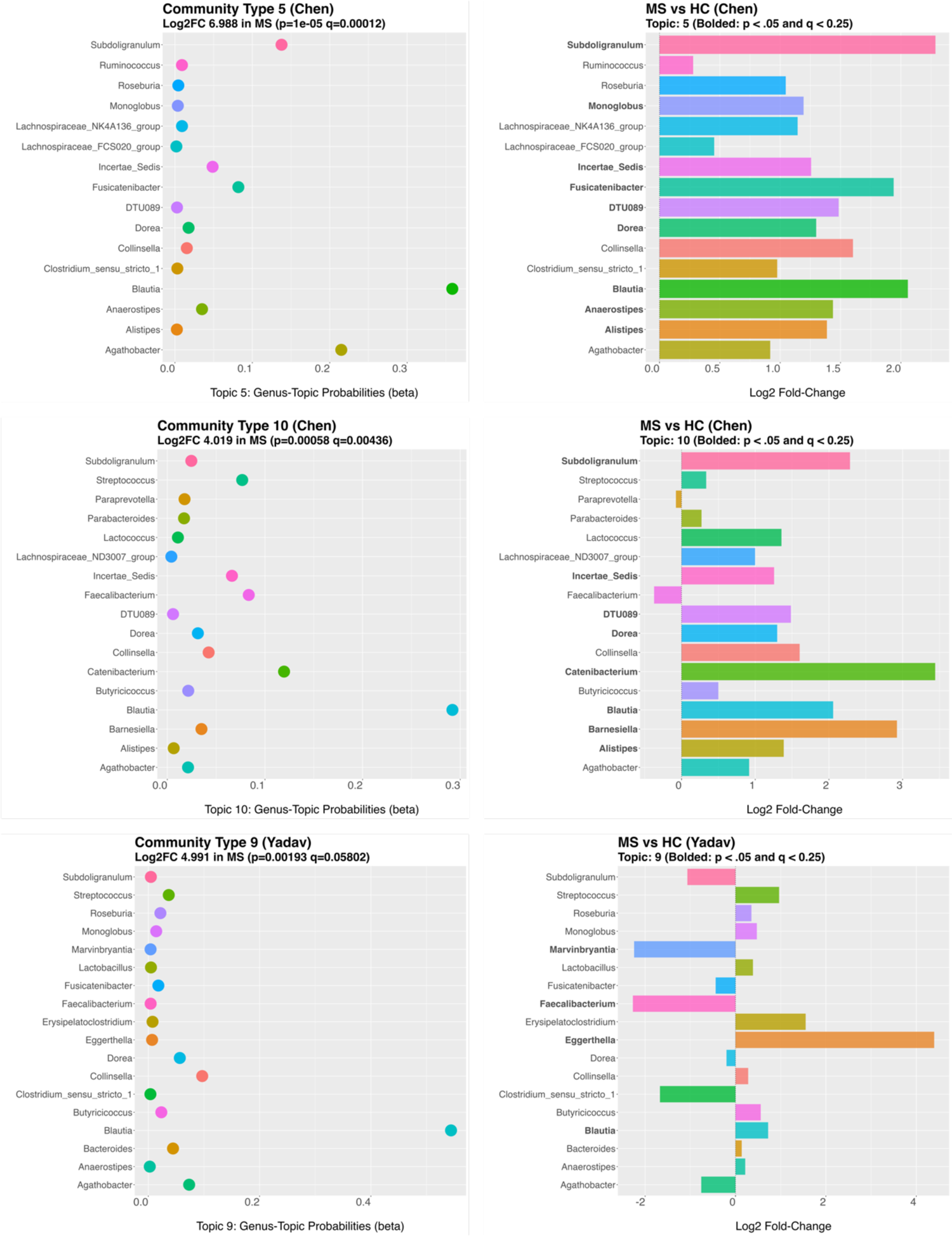
RRMS gut microbiome Community Type B structures found in Chen and validated in Yadav. Specifically, communities Chen Topic 5, Chen Topic 10, and Yadav Topic 9.

**Fig 4a:**
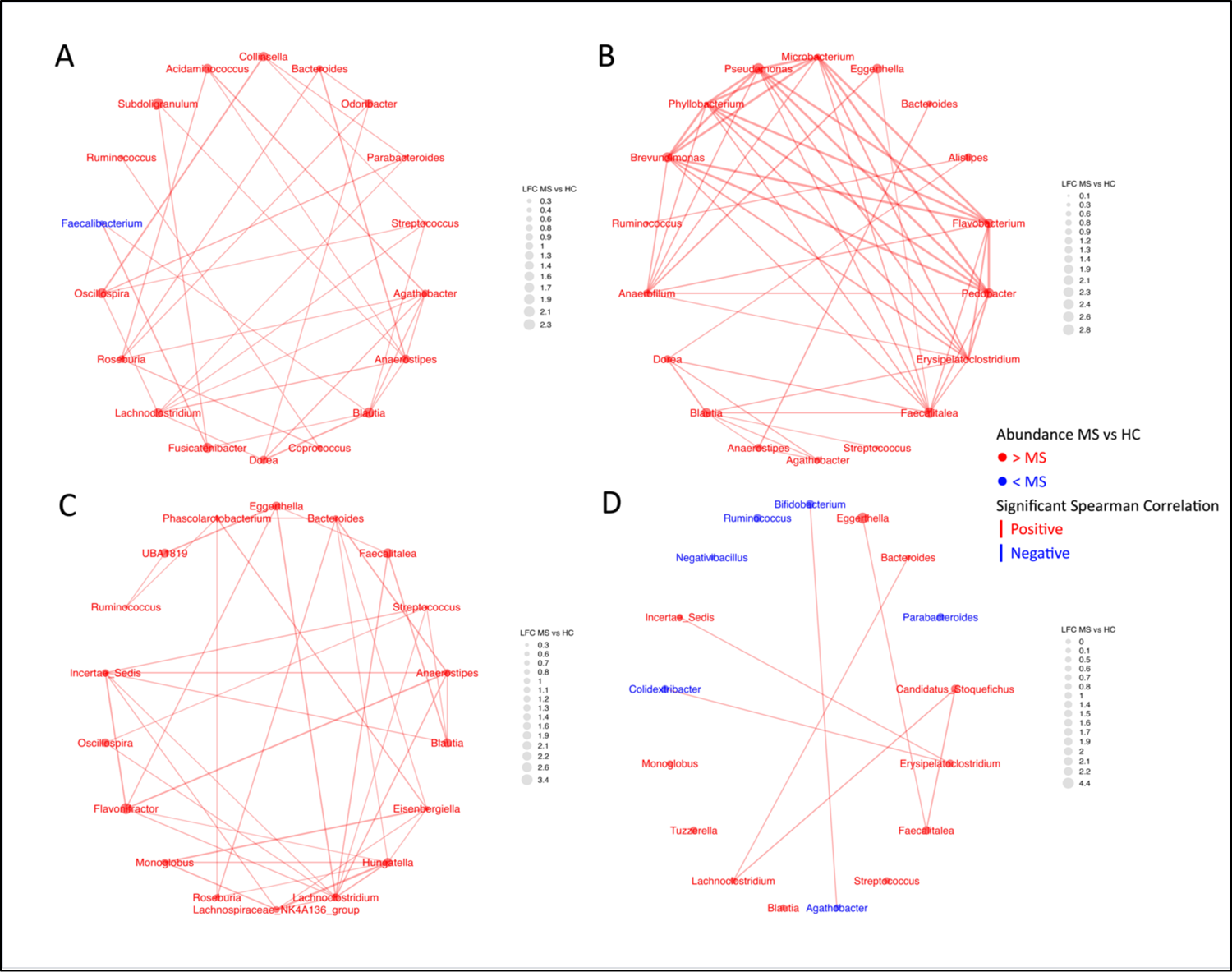
Network plots reveal the complexity of the topics making up Community Type A. Significant spearman correlations are represented by the edges. Thickness of edges corresponds to the strength of the correlation. A) Chen Topic 4; B) Chen Topic 6; C) Chen Topic 23; D) Yadav Topic 8.

**Fig 4b:**
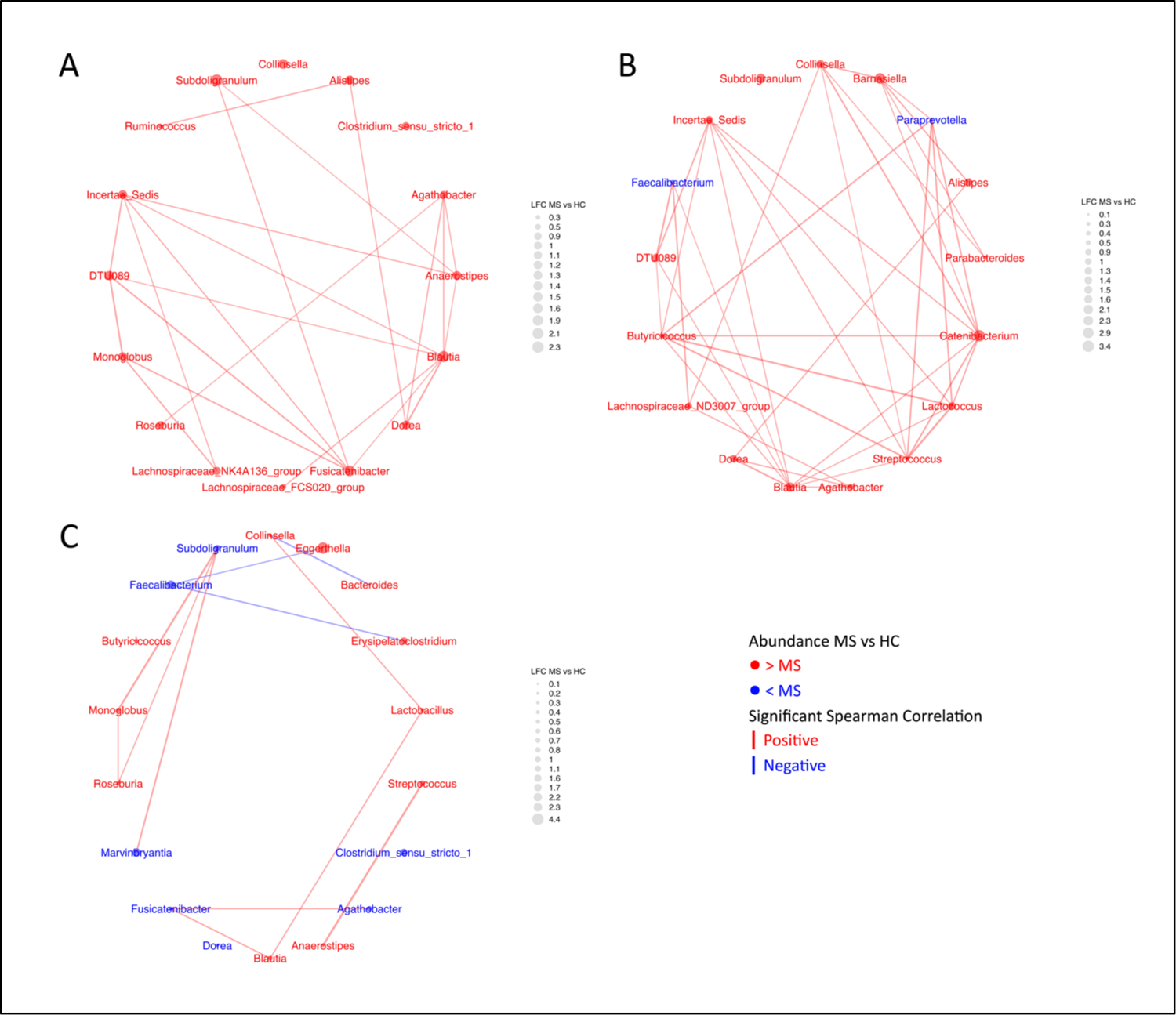
Network plots reveal the complexity of the topics making up Community Type B. Significant spearman correlations are represented by the edges. Thickness of edges corresponds to the strength of the correlation. A) Chen Topic 5; B) Chen Topic 10; C) Yadav Topic 9.

### Highly Assigned Genera and Their Directional Abundance in Validated Topics

We next examined the genera with high probabilities of assignment to these community types. In Community Type A, *Bacteroides* was the most often assigned genera, followed by *Blautia*, both were increased in RRMS compared to HC. Many other genera were also assigned to this community and were higher in RRMS than HC, including *Streptococcus, Eggerthella, Faecalitalea,* and *Lachnoclostridium*.

Several genera were assigned to Community Type A that varied in directional abundance between datasets. For example, *Ruminococcus* and *Agathobacter* were higher in the RRMS group in the Chen dataset and lower in the RRMS group in the Yadav dataset. Inversely, *Erysipelatoclostridium* was lower in RRMS in Chen and higher in RRMS in Yadav.

In Community Type B *Blautia* was the most often assigned genus and increased in RRMS compared to HC. Other genera enriched in RRMS and assigned to this community include *Dorea, Streptococcus*, *Butryicoccus, Roseburia, Monoglobus*, and *Anaerostipes.* The only genera depleted in RRMS across all the communities making up Community Type B was *Faecalibacterium.* Several other genera were important to Community Type B but varied in abundance between datasets. Specifically, *Subdoligranulum, Agathobacter, Clostridium sensu stricto 1, and Fusicatenibacter* were higher in RRMS patients in Chen, but lower in RRMS patients in Yadav.

### Differential Abundance Testing Within Topics

We assessed the assigned genera’s differential abundance within the statistically significant community types. Community Type A comprised many enriched genera; however, only *Eggerthella* and *Blautia* were significantly increased in RRMS compared to HC in both the Chen and Yadav datasets. None of the significantly depleted genera differed between RRMS and HC in abundance in both datasets. In Community Type B several genera were identified as differing in abundance, but the only shared significant finding between datasets was the increase in *Blautia* in RRMS.

## Discussion

We hypothesized that communities of microbes might be associated with RRMS patients when compared to healthy controls and that microbes not identified by one-at-a-time differential abundance testing approaches would be important to these dysbiotic community types. As such we utilized our previously published data and performed topic modeling on this dataset to look for community types associated with RRMS. Out of 30 topics assessed, we identified 10 that were more often associated with RRMS when compared to HC, and we validated these findings utilizing a separate dataset.

Several themes were found in Community Type A and Community Type B, suggesting similar dysbiotic communities associated with RRMS. We found that *Bacteroides* was one of the most often assigned genera. This genus was higher in RRMS than HC in Chen, and validated in Yadav, but did not reach statistical significance at this sample size in either dataset. This finding highlights the possibility that differences in clusters of microbes might be more important than differences in specific microbes in the dysbiotic gut microbiome communities seen in MS patients. *Blautia* also had a high assignment probability and was enriched in RRMS patients in both datasets. In multiple studies, *Blautia* has been linked to MS [17, 20]. Functionally its enrichment and depletion in the gut have both been linked to inflammatory diseases (enrichment: breast cancer [4], inflammatory bowel syndrome [21], and MS [10, 20]; depletion: Crohn’s disease [22], colorectal cancer [1], and MS [17]). Additionally, *Dorea* was highly assigned to these Community Types, and although *Dorea* is usually considered a gut commensal, its increased abundance has been linked to MS [20] and other inflammatory diseases such as Crohn’s disease [21]. Specifically, the pro-inflammatory effects of *Dorea* could be due to the fact that some species of Dorea can induce IFNy, metabolize sialic acids, and degrade mucin [23, 24]. Furthermore, *Blautia* utilizes gases produced by *Dorea* [25], thus these inter-bacterial interactions could be important to the gut microbiome community impact on RRMS patients.

*Eggerthella, Roseburia*, and *Anaerostipes*, were also assigned to these RRMS community topics and found to be higher in RRMS compared to HC. *Eggerthella* was identified in both datasets as being significantly higher. This increased abundance has been identified in multiple MS studies [12, 16, 20] and other autoimmune disorders like systemic lupus erythematosus [26]. Although we found *Anaerostipes* and *Roseburia* to be increased in RRMS, other studies have found the inverse [18, 20]. Even with these differences, this highlights that these genera are important to the RRMS community structure and future studies are needed to sort out their function and apparent abundance changes.

Several genera linked to gut permeability were assigned to the RRMS community types including *Streptococcus, Lachnoclostridium, Faecalibacterium,* and *Faecalitalea*. *Streptococcus* had a higher relative abundance in RRMS patients compared to HC, but again associations for this specific genus did not reach statistical significance. *Streptococcus* species have been shown to cross the epithelial barrier and translocate systemically [27], thus have the ability to induce systemic inflammation. They can also cross the blood-brain-barrier (BBB) [28, 29], which is of interest in MS research, as the gut-brain axis is often implicated in the pathobiology of this disease. One hypothesized mechanism of action is that inflammation of the intestinal barrier, potentially due to a lack of short-chain fatty acids (SCFAs) or other immunological changes, results in gut dysbiosis [30]. This allows pathogens and bacterial products to either affect the CNS directly through neuro-immune-endocrine pathways or indirectly by inducing systemic inflammation due to the translocation of bacteria and their products into the bloodstream and then to the CNS. As several microbes and microbial by-products have been identified in the CNS of MS patients [31], the gut-brain axis has gained traction and is important to consider when understanding the etiopathology of MS. Additionally, higher abundances of *Lachnoclostridium* have been linked to reduced levels of acetate [32]. Acetate is a SCFA that has been associated with a healthy gut microbiome and a developed immune system [33], reducing this protection would increase the penetration of the BBB in RRMS patients.

*Faecalibacterium* was also assigned to these community types and was lower in RRMS patients than HC. This genus is a butyrate-producer and linked to a decrease in intestinal inflammation [34]. Thus, along with the increase of several genera, a decrease in others such as *Faecalibacterium* are important to the community structure of the dysbiotic gut microbiome of RRMS patients and possibly gut permeability. Of note, *Faecalitalea* was assigned to the RRMS community types and was increased in RRMS patients compared to HC. This genus is thought to be beneficial as it can ferment many sugars and its major end product is also butyric acid [35]. Butyric acid is considered to support the integrity of the gut [36].

Collectively, our findings indicate that the complex dysbiotic microbiota in RRMS patients can be characterized by a diverse community of bacteria specifically comprising a reduction in beneficial symbiont bacteria, an increase in potentially harmful pathogenic bacteria, and an overall shift of certain commensal bacteria towards a pathobiont phenotype. As a number of bacteria in these communities don’t reach statistical significance on their own, our findings highlight that the collective impact of these bacteria is greater than their individual effect. Thus, a healthy or disease phenotype outcome can be attributed to the balance between symbionts and pathobionts shifting towards pathobionts. It seems there are certain keystone symbionts species, such as *Faecalibacterium,* which are mostly associated with a healthy phenotype, likely due to their inability to adjust to environmental changes lacking nutritional sources such as dietary fibers [37]. However, other commensal gut bacteria, such as *Bacteroides*, *Blautia,* and *Eggerthella spp.,* can be more flexible due to their adaptability to thrive in diverse conditions and utilize a wide range of substrates as a food source. They can efficiently switch their metabolic pathways and enzymatic activities to utilize different nutrients, ensuring their survival and maintenance in the ever-changing gut ecosystem [38, 39]. Based on our data, we propose a potential mechanism through which healthy gut microbiota can be converted to a dysbiotic phenotype. This can be explained through a term used in social sciences: “bottom-up influencer”, where peripheral pressure leads to changes in a central authority [40]. In a steady state, the gut microbiota is dominated by keystone species such as *Faecalibacterium* which had been shown to be highly abundant in human populations. These keystone species regulate the properties of other peripheral members of this community (e.g., *Blautia, Dorea,* and *Eggerthella)* to sustain a healthy state by producing beneficial metabolites required for maintaining an intact gut barrier and inducing anti-inflammatory responses. However, in certain scenarios such as infection, dietary, or environmental changes, peripheral members can adjust to the changing environment better than the keystone species resulting in their higher abundance due to the reduction/depletion of keystone species. A higher abundance of certain commensal bacteria in a disease state suggests these community members might acquire an inflammatory potential in the absence of keystone species. As there is lots of heterogeneity in human populations, certain individuals might be more prone to these bottom-up influencers, thus being more likely to have a dysbiotic phenotype and further, a predisposition to diseases such as MS. It is possible that some of these bacteria start metabolizing or degrading host-derived nutrients such as mucins or synthesize immunostimulatory LPS as shown by us recently [41]. This can result in a compromised gut barrier, “leaky gut”, and translocation of bacteria or bacterial by-products into circulation leading to systemic inflammation. Finally, this transition provides a conducive environment for opportunistic bacteria to colonize and perpetuate a pro-inflammatory response.

However, there are a number of unknowns, such as what are the factors promoting dysbiosis, why certain individuals are more prone than others and most importantly, whether dysbiosis can be corrected through diet (prebiotic) or microbiota replacement (probiotic) or both (symbiotic). Thus, future studies are warranted to determine these factors to better understand the mechanism promoting dysbiosis. This understanding would help harness the enormous potential of the gut microbiome as a future diagnostic and therapeutic agent.

Our findings here are in line with many prior findings on the dysbiotic gut microbiome of RRMS patients. In addition, with the use of topic modeling, we observed associations for community structures related to RRMS that cannot be identified with differential abundance testing. These findings should be further validated with more datasets and diverse cohorts but highlight the potential of topic modeling in microbiome research. In the future, we hope that topic modeling will be incorporated with traditional statistical approaches for microbiome analysis and help provide a better picture of the microbiome as a whole in complex diseases such as RRMS.

## Methods

### Clinical and Sequence Data

The clinical and 16s sequence data for our exploratory analysis were obtained from Chen et. al. 2016 [11], a prior publication from our group. The clinical and 16s sequence data for our validation analysis were obtained from Yadav et. al. 2022 [3], a separate publication from our group. For simplicity, each dataset is referred to using the first author’s last name (i.e., Chen and Yadav).

### 16s Sequence Data Processing

Sequence data for the V3-V4 region of the bacterial 16s rRNA gene for each study was obtained from the National Center for Biotechnology Information (NCBI) Sequence Read Archive (SRA) under the BioProject numbers PRJNA335855 and PRJNA732670. The demographic data for all datasets can be found in Supplementary Table 1.

The sequence data was downloaded utilizing the SRA toolkit, denoised with DADA2 [42] using the default parameters and trimming of the forward and reverse reads at 240 and 180nt, respectively, for Chen, and trimming of the forward and reverse reads at 290 and 275nt, respectively, for Yadav. The taxonomy was assigned using the assignTaxonomy function and the Silva database (Version 138). Low prevalence features (relative abundance < 1e-5) were removed. Post-filtering, the reads were aggregated at the genus level for analysis.

### Statistical Analysis and Topic Modeling

Statistical analyses were performed using the R environment for statistical computing and graphics (version 4.2.3). We first built a phyloseq [43] object using the genus abundance table (i.e., genus-level phylotypes) and metadata to facilitate the statistical analysis.

We utilized the FindTopicNumber function from the ldatuning [44] package to identify an optimal latent topic number for our model based on the CaoJuan2009 [45] and Arun2010 [46] metrics. The method = “VEM” option was selected to perform variational inference when deriving the latent topics. This was performed on each dataset and an average of the ideal topic numbers was selected. A total of 30 topics was chosen based on this approach (Supplementary Figure 1).

To derive the final set of topics we used the LDA function from the topicmodels [47] package to perform the latent Dirichlet allocation on each dataset. This model was chosen as it allows for fractional membership, or the allowance of assignment to multiple topics, when assigning reads to the underlying community types. We then extracted the beta and gamma probability matrices from our topic model using tidytext package [48] and multiplied the per-document-per-topic probabilities by the read count for each sample to assign reads to each topic (i.e., generate a document-term matrix). A new phyloseq object was then built with the document-term matrix serving as the abundance table.

### Validation of Topics Found in Original Dataset

We extracted the topic-term-probability matrix from the Chen and Yadav LDA models and assessed the similarity in the community structure (topics) between our exploratory and validation data by calculating the cosine similarity matrix for each topic. The communities that had a cosine similarity of 0.80 or higher were considered to reflect similar community types identified independently in each dataset, and thus validating the findings from the Chen data.

### Differential Abundance of Topics and Bacteria within Topics

To assess differences in the relative abundance of each community type between samples collected from RRMS patients and HC we performed a differential abundance analysis using the LinDA (linear models for differential abundance analysis) function from the MicrobiomeStat [49] package with feature.dat.type = “count” and is.winsor = F for community type comparison, and is.winsor = T for genera comparison. The Benjamini-Hochberg false discovery rate (FDR) correction was applied to account for the multiple testing. LinDA was also used to test for differences in the genus-level relative abundances. Community Types (topics) and bacteria with a p ≤ 0.05 and a q-value ≤ 0.25 were considered differentially abundant.

## Data Availability

The sequence data used for analysis can be found at the National Center for Biotechnology Information (NCBI) Sequence Read Archive (SRA) under the BioProject numbers PRJNA335855 and PRJNA732670. On GitHub (https://github.com/raeshrode/TheFullPicture_Article) are the following data and scripts: (1) R script to analyze Chen data, Yadav data, and cosine similarity; (2) R environments after topic model analysis of Chen and Yadav datasets, (3) abundance tables, metadata, and taxa tables for each dataset, and (4) the R scripts to create the network plots.

## Author Contributions

RLS, AKM, and NJO contributed to the study conception and design. Data collection, processing, and analysis were performed by RLS. NJO provided guidance in data analysis. The manuscript was written by RLS, AKM, and NJO. All authors read and approved the final manuscript.

## Supporting Information

Supplementary Figure 1: The ideal topic number for each dataset is at the minimum value that both metrics generally reach. For the Chen dataset 33 topics was ideal and for Yadav 27 topics was ideal. On average, 30 topics is ideal for these datasets and was selected.

Supplementary Figure 2: a) The 10 of 30 significant topics found in the Chen dataset. b) The 4 of 30 significant topics found in the Yadav dataset. The left-hand side contains the probability of the genus being assigned to the topic. The right-hand side contains the abundance comparison between MS and HC of the highly assigned bacteria. Bolded genera are significant. Positive values indicate a higher abundance in MS compared to HC.

Supplementary Table 1: Demographics for all datasets.

